# Cisternostomy facilitates clearance of metabolic waste from cerebrospinal fluid in patients with traumatic brain injury

**DOI:** 10.1101/2025.01.22.634226

**Authors:** Weiwei He, Shuai Gao, Lulu Du, Tangrui Han, Di Yao, Hao Wu, Qiang Li, Yonghong Wang, Woo-ping Ge

## Abstract

Decompressive craniotomy, a common intervention for traumatic brain injury (TBI), can fail to effectively alleviate patient symptoms. Cisternostomy, reported for cistern drainage in TBI patients, has shown efficacy in reducing intracranial pressure and clearing detritus resulting from brain hemorrhage. However, the mechanisms underlying its effectiveness remain largely unknown. Here, we utilized non-targeted metabolomics to analyze cerebrospinal fluid from cisterns alongside peripheral blood samples from TBI patients undergoing cisternostomy. Through a systematic comparison of the cisternal cerebrospinal fluid and blood plasma metabolomes, we identified multiple blood-enriched metabolites, including betaine, triethanolamine, and proline, that were efficiently cleared during the acute stage of TBI. Notably, two metabolites linked to arginine metabolism and the urea cycle, N8-acetylspermidine and N-acetylputrescine, showed significant reductions that correlated with improvements in the Glasgow Coma Scale. Our findings indicate that cisternostomy effectively removes blood-derived substances and aids the recovery of patients with acute-stage TBI.

## Introduction

The brain vasculature is uniquely characterized by the protective blood-brain barrier (BBB), which serves as the critical interface between the bloodstream and central nervous system (CNS). The BBB selectively filters metabolites between the blood and cerebrospinal fluid (CSF), and this mechanism is essential for maintaining CNS homeostasis^1^. The CSF is predominantly produced by the choroid plexus, and it circulates throughout the CNS and is subsequently drained via arachnoid granulations and the glymphatic system^1, 2^. Substances that penetrate the BBB and reach brain tissue help regulate both the physical and chemical environment of the CNS; hence, any alteration or disruption of the BBB may expose the brain to harmful substances from the blood, which can lead to neuronal and glial dysfunction and ultimately cause brain damage. Thus, controlling the exchange of substances between the blood and the brain parenchyma is essential for brain function and protection^3, 4, 5^.

Traumatic brain injury (TBI) is a major cause of disability and death worldwide, affecting an estimated 50 million people annually^6^ ^7^ ^8^. TBI can cause diffuse brain injury, which typically results from exposure of the head to a sudden accelerating or decelerating force. This can lead to a spectrum of pathological outcomes, including axonal injury, cerebral swelling, contusions, lacerations, and intracranial hemorrhage^9,10,11,12^. TBI is often accompanied by disruption of the BBB, thus impairing the transport of essential nutrients from the blood to the brain parenchyma^13^. Additionally, any compromise in BBB may facilitate the infiltration of blood-derived toxins, inflammatory molecules, and immune cells into the brain tissue^14 15 16 17^. This breach can trigger extensive neuroinflammation, oxidative stress, and ultimately neuronal damage^18, 19^. Consequently, BBB disruption or damage to the vasculature may lead to the development of brain edema, elevated intracranial pressure, or impaired cerebral autoregulation^15, 16, 20^. These pathological changes interfere with normal neuronal function and connectivity, ultimately resulting in diverse forms of brain dysfunction including cognitive deficits^16, 21^, motor impairments^22, 23^, and increased susceptibility to neurodegenerative diseases^24^ ^25^ ^26^ ^27^ ^28^. Therefore, reestablishing the integrity of the BBB as soon as possible after TBI would restore its ability to clear potentially noxious or exogenous substances from the brain, help prevent secondary brain injury, and promote recovery^1^.

Decompressive craniectomy is a commonly performed neurosurgical procedure aimed at reducing intracranial pressure in patients with TBI^29^. Although it relieves the pressure by creating physical space in which the brain can expand, this intervention does not accelerate the removal of blood-derived substances that infiltrate the CSF post-injury^30^. Perhaps for this reason, decompressive craniectomy has not consistently shown efficacy in treating cerebral edema, improving CSF circulation, or enhancing glymphatic system clearance^31^. Furthermore, due to the risks associated with bone flap loss, many patients require cranioplasty after the initial procedure^32^. Therefore, there is an urgent need to develop novel treatment modalities for TBI^30^.

Since the cisternostomy procedure was reported^33, 34, 35^, it is now used in the acute phase of severe TBI to mitigate intracranial pressure by creating a stoma in the intracranial basal cisterns, facilitating CSF drainage^36–39, 40^. This approach aims to prevent edema and thus secondary brain injury^20,41,42^. However, the widespread adoption of cisternostomy has been limited, primarily due to uncertainties regarding the relationship between the efficacy of the procedure and the underlying mechanisms^40, 43, 44^. Additionally, the ability of the procedure to remove harmful substances (including metabolites) from the CSF—particularly blood-derived substances following cerebral hemorrhage—remains uncertain^45^.

To address the challenges limiting the widespread use of cisternostomy for TBI and evaluate its efficacy, we performed a systematic metabolomic analysis of CSF and blood plasma samples (acquired 5–8 days post-injury) from patients who experienced severe TBI. Our results demonstrate that cisternostomy, when administered during the acute stage after TBI, significantly reduced the levels of substances including metabolites from CSF and those enriched in blood. These findings provide evidence that cisternostomy provides substantial clinical benefit for managing TBI.

## Results

### Characterizing the difference in metabolic profiles between blood and clear cisternal CSF

To obtain and compare the metabolomes of blood and clear cisternal CSF from acute-stage TBI patients, peripheral blood plasma (n = 20 samples) and clear CSF (n = 20 samples; clear cisternal was defined as cisternal CSF from patients with high Glasgow Coma Scale (GCS) scores from 12–15 and without hemolysis) were collected for non-targeted metabolomic profiling (**Fig. 1a**). Overall, there were substantial differences between the metabolomes of blood and clear CSF (**Fig. 1b**). A total of 2,159 metabolic features (briefly, we used metabolites in subsequent text) were observed to be enriched in blood, whereas 4,164 were enriched in clear CSF (**Fig. 1c**, filtered by fold change, FC, of >1.5 and an adjusted *p* < 0.05); these accounted for 16.46% and 31.78%, respectively, of the 13,114 metabolites we measured (**Fig. 1d**). Subsequent analysis of the chemical species among these enriched metabolites revealed a prevalence of lipids, particularly various glycerophospholipids, in blood. In contrast, water-soluble metabolites were relatively enriched in clear cisternal CSF (**Fig. 1e**). We next conducted a biochemical pathway analysis of the enriched metabolites and found that the linoleic acid pathway was specifically enriched in blood (**Supplementary Fig. 1a**). Additionally, pathways associated with amino acids, coenzymes, and vitamins (e.g., nicotinate and nicotinamide metabolism pathway) were found to be enriched in cisternal CSF (**Supplementary Fig. 1b**). We further performed a KEGG pathway enrichment analysis of the metabolites enriched in blood and CSF (**Supplementary Fig. 2**). A substantial number of metabolic pathways were significantly enriched in blood, including lipid metabolism–related pathways such as choline and glycerophospholipids. In contrast, certain other subclasses of metabolites were enriched in cisternal CSF, including pathways related to the biosynthesis of tyrosine, arginine and proline and the histidine metabolism pathway (**Supplementary Fig. 2**).

**Fig. 1.**
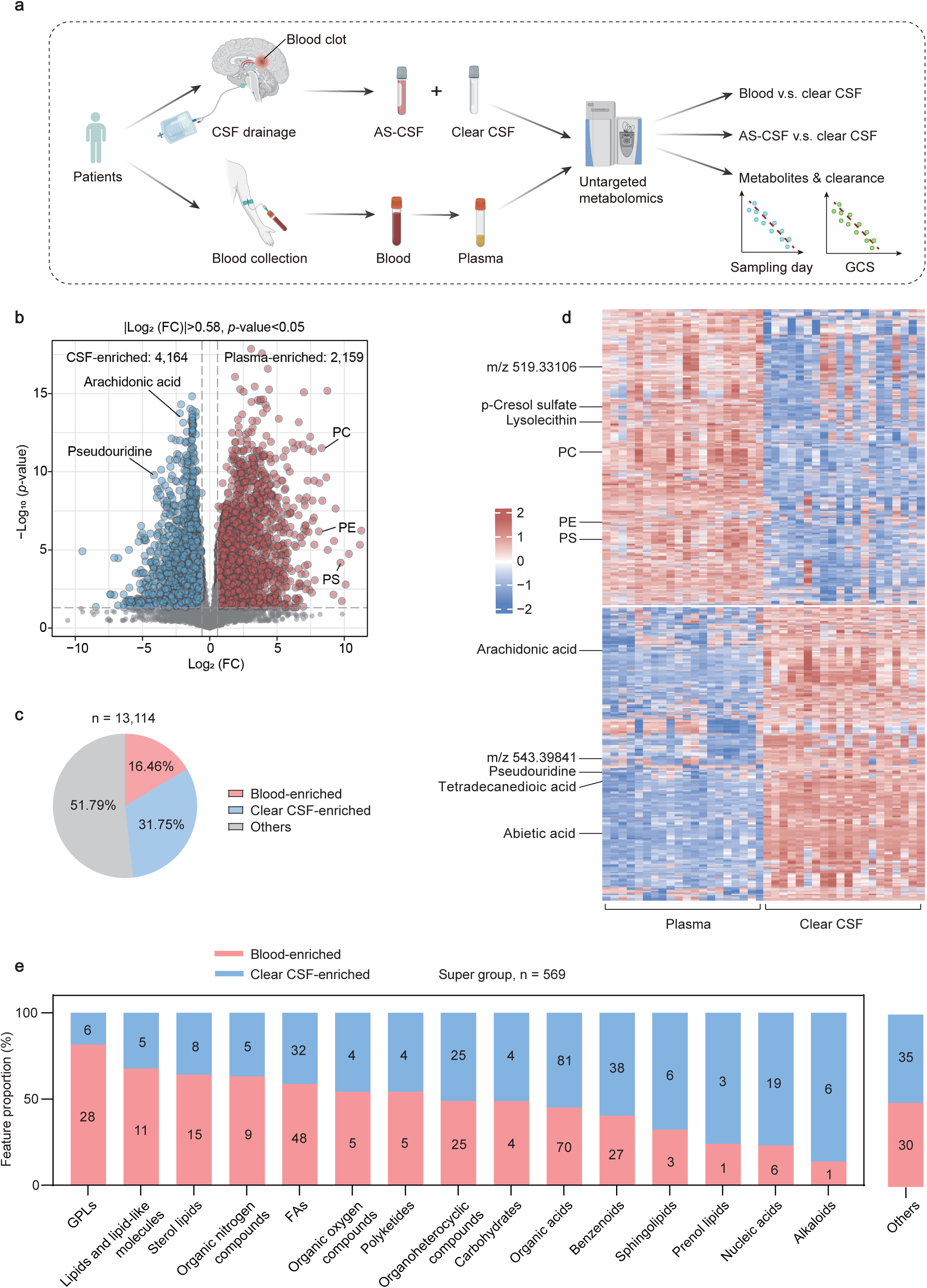
Comparison of metabolomes of peripheral plasma and clear cisternal cerebrospinal fluid (CSF) from patients with TBI. **a)**, Schematic illustration demonstrating the strategy for collecting plasma and CSF samples, followed by metabolomic measurements and analysis. **b)**, Volcano plot showing the most significantly different metabolic features between plasma and cisternal CSF (p < 0.05, |log₂FC| > 0.58, i.e., FC > 1.5 or FC < 0.67). *FC*: fold change. **c)**, Percentage of differential metabolites between plasma and CSF samples among all metabolites measured. **d)**, Heatmap of the top 3,000 differential metabolic features between plasma and clear CSF (n = xx plasma samples, n = xx CSF samples). Data were obtained using non-targeted metabolomic analysis with LC-MS (see *Methods* for details). The color scale represents the levels of standardized metabolites. **e)**, Enrichment analysis of metabolites from different sub-clusters. The metabolite clusters were categorized based on chemical structures from differential and annotated blood-enriched and cisternal CSF–enriched metabolites. A total of 569 annotated metabolites were identified among all differential metabolites.

To further analyze the chemical species represented by the metabolites enriched in CSF and blood plasma, we compared their retention time (RT, via liquid chromatography), molecular polarity, and the calculated molecular mass (Cal. MW) across groups. Notably, metabolites that were abundant in blood were concentrated within two distinct molecular weight ranges, namely 50–200 Da and 750–850 Da, with corresponding RTs within the 0–2 min timeframe (RTs obtained from the HILIC chromatographic column). This observation suggested that these metabolites may represent macromolecular lipids with low polarity based on their corresponding fragment. KEGG pathway enrichment analysis was performed on the top 20% of metabolites (430 in total, comprising 160 annotated metabolites), revealing that these metabolites were mainly glycerophospholipids, linoleic acids, unsaturated fatty acids, and other lipid-related metabolites (**Supplementary Fig. 3**). The highly abundant metabolites in CSF were observed to cluster within the molecular weight ranges of 50–300 Da and 500–650 Da, with RTs predominantly between 3 and 5 min. This suggested these metabolites represented small molecules with high polarity. KEGG pathway enrichment analysis conducted on the top 20% of metabolites (comprising 832 metabolites in total, with approximately 170 annotated metabolites) indicated that the predominant metabolites in clear CSF were small-molecule metabolites of histidine, tyrosine, and nicotinoids (**Supplementary Fig. 4**). These results demonstrated that significant differences existed between the metabolomes of plasma and cisternal CSF, with each displaying distinct chemical characteristics.

### Characterizing the differences in molecular species between clear CSF and cisternal CSF in samples from acute-stage TBI patients

To elucidate the molecular mechanisms by which cisternostomy contributes to the recovery of TBI patients, we conducted a comparative metabolomic analysis of CSF collected at two different stages of recovery. Specifically, we examined the differences between cisternal CSF obtained during the acute stage of TBI (AS-CSF, n=16 samples from 5 subjects) and clear CSF collected during the late stage (n=20 samples from 6 subjects) from TBI patients who underwent cisternostomy. Significant differences were observed between the metabolomes of AS-CSF and clear CSF (**Fig. 2a**, FC > 1.5, adjusted *p* < 0.05). Of the 13,114 metabolites analyzed, 10.49% (1,376 metabolites) were enriched in AS-CSF, whereas 13.07% (1,714 metabolites) were enriched in clear CSF (**Fig. 2b, c**).

**Fig. 2.**
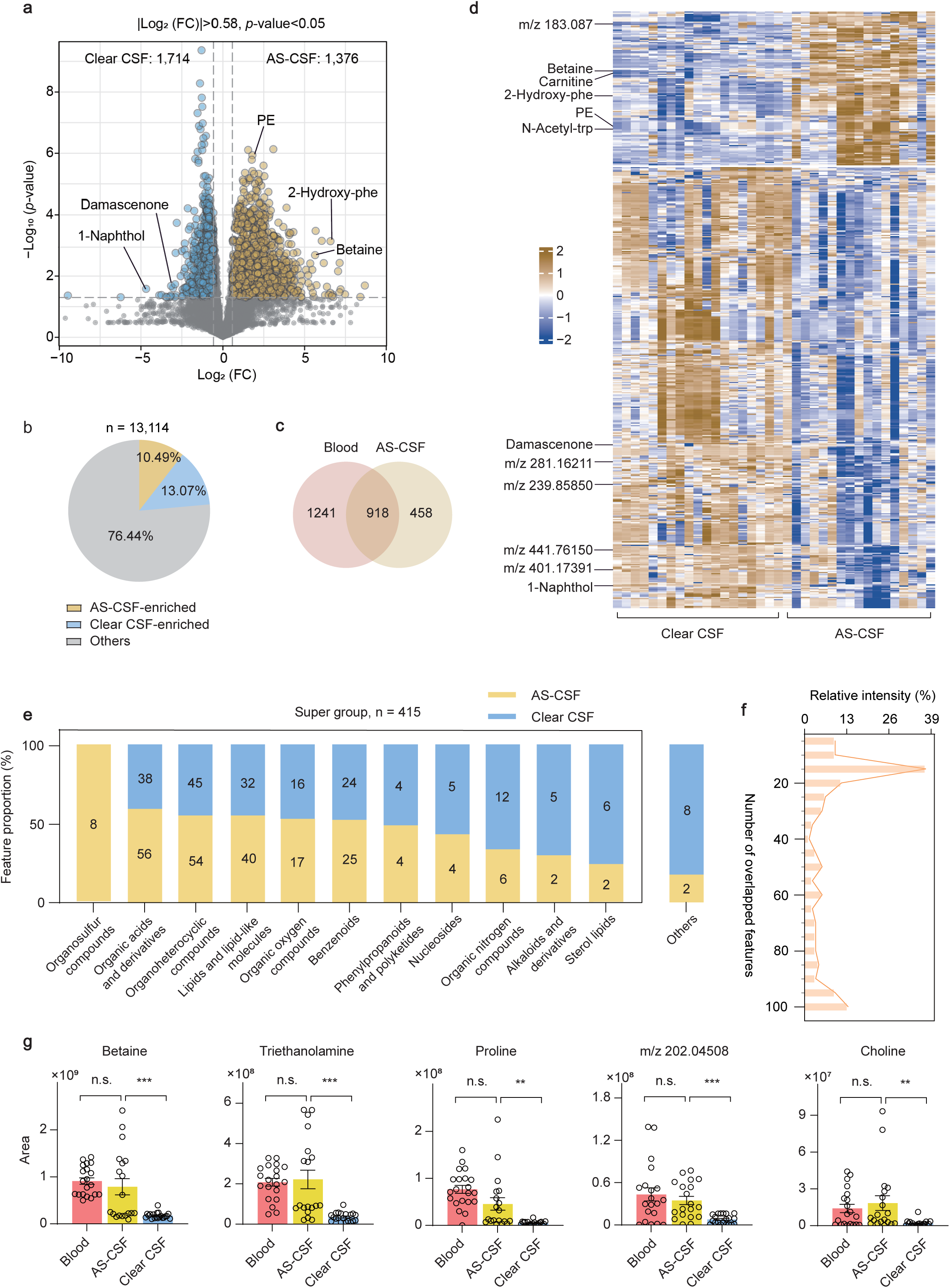
Metabolomic comparison of acute-stage cisternal CSF (AS-CSF) and clear cisternal CSF. **a)**, Heatmap of the top 3,000 differential metabolic features between AS-CSF and clear CSF. The color scale represents the levels of standardized metabolites. **b)**, Volcano plot showing the most significantly different features and corresponding metabolites between AS-CSF and clear CSF (p < 0.05, |log₂FC| > 0.58, i.e., FC > 1.5 or FC < 0.67). **c)**, Percentage of metabolites for which differences were statistically significant between AS-CSF and clear CSF. **d)**, Venn diagram illustrating the overlap between blood-enriched and AS-CSF-enriched metabolic features, with 918 overlapping features identified. **e)**, Enrichment analysis of metabolites based on the chemical structures of metabolites that were enriched in AS-CSF or clear CSF. A total of 415 annotated metabolites were divided into 12 distinct groups. **f)**, Proportion of AS-CSF-enriched features within blood-enriched features, categorized based on the ranking of relative intensity. **g)**, Intensity of five representative metabolites (betaine, triethanolamine, proline, *m/z* 202/04508, and choline) which were enriched in blood and AS-CSF but significantly reduced in clear CSF.

To investigate the proportion of blood-derived metabolites present in AS-CSF, we integrated the blood-enriched metabolites identified earlier with the metabolites enriched in AS-CSF (**Fig. 1**). Two-thirds of the metabolites (n = 918) enriched in AS-CSF overlapped with the blood-enriched metabolites (**Fig. 2d**), suggesting that a substantial portion of the metabolites in AS-CSF may have originated from blood clots or were a consequence of cerebral hemorrhage. Among the blood-derived metabolites, lipophilic substances were more prevalent in AS-CSF, whereas hydrophilic substances were relatively enriched in clear CSF (**Supplementary Fig. 5** and **Fig. 2e**). The above observations revealed similar enrichment characteristics to the above differential metabolites between blood and clear CSF (**Fig. 1e**).

To determine the likelihood that blood-enriched metabolites were discharged into AS-CSF, the 2,159 blood-enriched metabolites (**Fig. 1c**) were divided into 20 equivalent subgroups, i.e., each group contained 108 (∼5%) metabolites. These subgroups were plotted in order of descending relative abundance. Simultaneously, the 918 metabolites that were enriched in both blood and AS-CSF were also ranked in descending order based on abundance in AS-CSF and then divided into 20 subgroups of 46 metabolites (5%) each. We then compared the distribution of these 46 AS-CSF-enriched metabolites with the 108 blood metabolites in the corresponding ordinal section. Intriguingly, this analysis revealed a correlation between decreased abundance of a metabolite in AS-CSF and its decreased metabolite in blood (**Fig. 2f**). Moreover, 57% of potential blood-derived metabolites in AS-CSF were concentrated in the first 6 sections, representing the top 30% of blood-enriched metabolites (**Fig. 2f**). These results indicated that these metabolites, which were enriched in AS-CSF compared with clear CSF, were likely derived from blood clots or leaked blood. The results also indicated that cisternal CSF drainage during cisternostomy led to the gradual discharge of abundant blood-derived substances.

To further assess the overlapping metabolites among clear CSF, AS-CSF, and blood, we compared the abundance of 88 blood-enriched metabolites and examined their abundance in AS-CSF. These blood-enriched metabolites could be categorized into three distinct subgroups (**Supplementary Fig. 6**). The first group comprised 49 metabolites that were most abundant in blood and least abundant in clear CSF. This group accounted for 55.68% of the 88 total blood-enriched metabolites, including betaine, proline, and the metabolite with m/z 220.04508. Within this group, certain metabolites showed a smaller disparity in abundance between AS-CSF and clear CSF. The second group comprised 21 members (23.86% of the total 88) including lysine, proline and valine (**Fig. 2g**) and was characterized by having the highest abundance in blood but also having high abundance in AS-CSF. The third group comprised 18 members characterized by having the highest abundance in AS-CSF including triethanolamine and choline, constituting 20.45% of the total (**Fig. 2g**), indicating that certain blood-derived metabolites may have become more concentrated upon entry into CSF due to its limited volume. The second and third groups of metabolites likely originated from blood and entered the CSF as a result of TBI-induced hemorrhage. The second group, in particular, likely originated from blood but became more concentrated upon entering the cisternal CSF due to the limited ventricular space.

### Identifying metabolites cleared from the brain during cisternostomy

To further investigate the efficacy of cisternostomy for clearing blood-derived metabolites from AS-CSF following TBI, we conducted a correlation analysis between GCS score, the timing of CSF sample collection from cistern drainage, and the abundance of 13,114 metabolites obtained from non-targeted metabolomic measurements (**Fig. 3a, b**). A significant number of metabolites correlated positively with sampling time and/or GCS score, representing 19.03% (2,496 metabolites) and 5.72% (598 metabolites), respectively, of the total 13,114 metabolites. Conversely, some metabolites correlated negatively with sampling time and/or GCS score, accounting for 4.56% (750 metabolites) and 2.13% (279 metabolites), respectively, of the total (**Fig. 3a, b**).

**Fig. 3.**
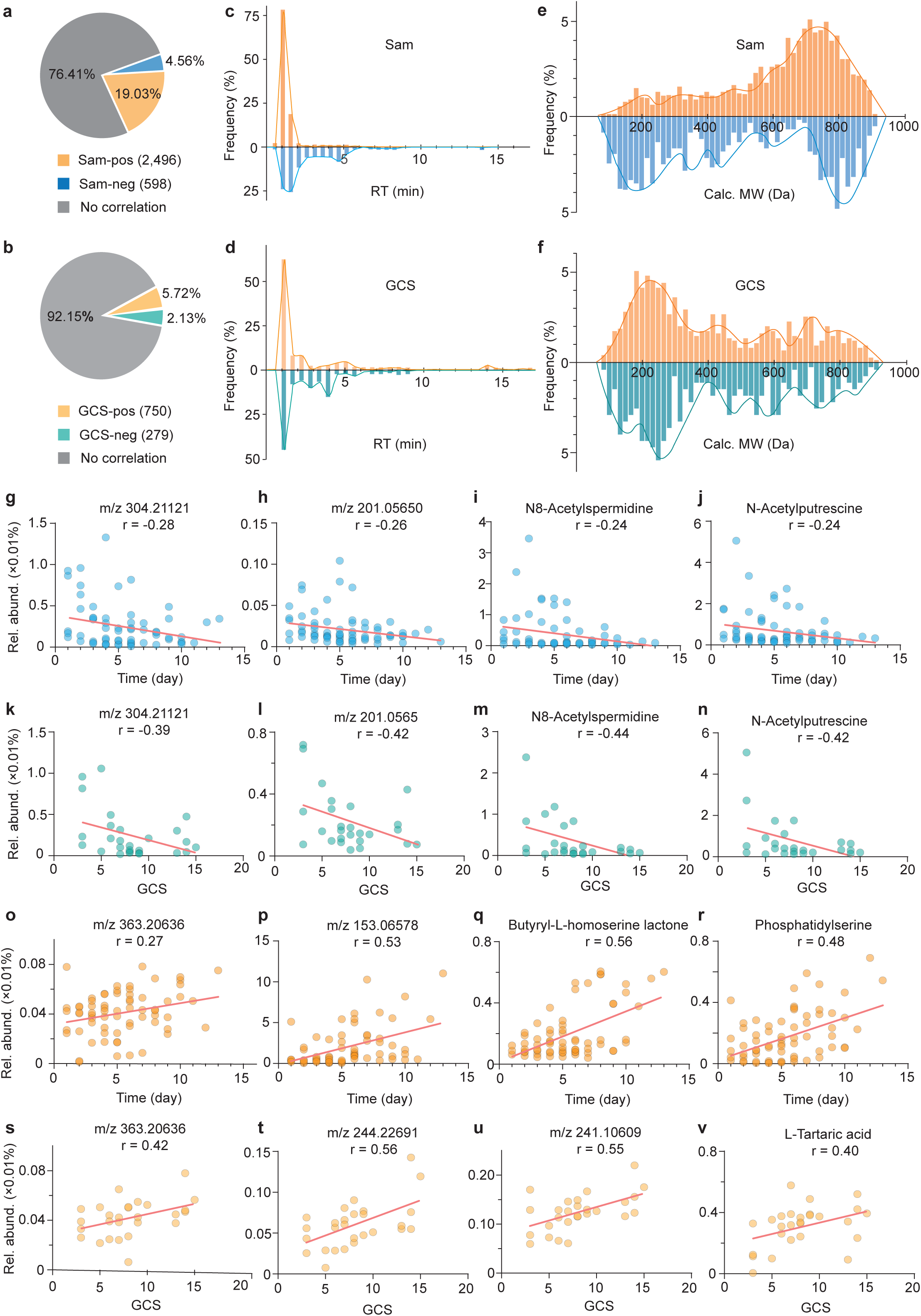
Identification of metabolites correlated with functional brain recovery in TBI patients. **a)**, Correlation analysis between cisternal metabolites and sampling day. Positively correlated metabolites (Sam-up, *r* > 0.35, *p* < 0.05) are shown in orange, accounting for 19.03% of the total (2,496 features). Negatively correlated metabolites (Sam-down, *r* < –0.35, *p* < 0.05) are shown in blue, accounting for 4.56% of the total (598 features). The remaining 76.41% of features exhibited no significant correlation. **b)**, Correlation analysis between cisternal metabolites and GCS score. Positively correlated metabolites (GCS-up, *r* > 0.35, *p* < 0.05) are shown in yellow, accounting for 5.72% of the total (750 features). Negatively correlated metabolites (GCS-down, *r* < –0.35, *p* < 0.05) are shown in green, accounting for 2.13% of the total (279 features). The remaining 92.15% of features exhibited no significant correlation. **c, d),** Distribution plots of retention time (RT, liquid chromatography) of metabolites correlated with sampling day (*Sam-up* and *Sam-down*) and GCS score (*GCS-up* and *GCS-down*). **e, f),** Distribution plots of the relative molecular weights of metabolites correlated with sampling day and GCS score. **g–j),** Representative metabolites that correlated negatively with sampling day, including *m/z* 304.21121, *m/z* 201.05650, N8-acetylspermidine, and N-acetylputrescine. **k–n)**, Representative metabolites that correlated negatively with GCS score, including *m/z* 304.21121, *m/z* 201.05650, N8-acetylspermidine, and N-acetylputrescine. **r)**, Representative metabolites that correlated positively with sampling day, including *m/z* 363.20636, *m/z* 153.06578, butyryl-L-homoserine lactone, and phosphatidylserine. s–v. Representative metabolites that correlated positively with GCS score, including *m/z* 363.20636, *m/z* 153.06578, butyryl-L-homoserine lactone, and phosphatidylserine.

To further elucidate the characteristics of the group of metabolites for which abundance correlated with cisternal CSF drainage (i.e., sampling time progression) and patients’ recovery (i.e., GCS score improvement), we analyzed their chemical properties. Specifically, we examined the distribution of RT (**Fig. 3c, d**) and molecular weight (**Fig. 3e, f**). The chemical properties of substances that correlated positively with sampling time and GCS (**Fig. 3a, b**) differed significantly from the properties that correlated negatively, particularly with respect to the RT distribution trends (**Fig. 3c, d**). The RTs of substances that correlated positively with sampling time and GCS score (Sam-pos and GCS-pos in **Fig. 3a, b**) were generally short, predominantly 0–3 min. Given that the HILIC chromatographic column has a polar stationary phase, strongly hydrophilic compounds had relatively longer RTs, suggesting that the positively correlated substances were more lipophilic. However, the overall RT distribution of the negatively correlated substances was relatively uniform, primarily 0–6 min, suggesting that the negatively correlated substances were more hydrophilic.

The molecular weight distribution trends differed slightly from trends seen in the analyses of sampling time (**Fig. 3e**) and GCS score (**Fig. 3f**). Specifically, substances that correlated positively with sampling time were predominantly 600–800 Da, whereas substances that correlated negatively were mainly <600 Da. These results suggested that the substances that correlated positively had larger molecular weights, whereas those that correlated negatively had smaller molecular weights. Nevertheless, the overall molecular weight distribution did not differ significantly between substances that correlated positively or negatively with GCS score, with the majority of these substances falling within the range 100–400 Da.

We further observed that the abundances of specific metabolites consistently correlated with the increase of GCS score and sampling time. For instance, the abundances of m/z 304.21121, m/z 201.05650, N8-acetylspermidine and N-acetylputrescine correlated negatively with GCS score and sampling time (**Fig. 3g–n**). This suggested that the concentrations of these metabolites progressively decreased in conjunction with cisternal CSF drainage and recovery from TBI. Conversely, the abundance of m/z 363.20636 correlated positively with both GCS score and sampling time, suggesting that its concentration progressively increased during drainage and recovery (**Fig. 3o, s**). We also found that the abundances of certain metabolites, including m/z 153.06578, butyryl-L-homoserine lactone and phosphatidylserine, were solely positively correlated with sampling time (**Fig. 3p–r**). In contrast, the abundances of m/z 244.22691, m/z 241.10609 and L-tartaric acid were exclusively positively correlated with GCS score (**Fig. 3t–v**).

### Clearance of metabolites by cisternostomy and its contribution to recovery

To determine whether cisternal CSF drainage during cisternostomy helps eliminate metabolic waste from the CSF of TBI patients, we analyzed the correlation between the abundance of specific metabolites in cisternal CSF and GCS score. First, we identified a pool of metabolites that correlated negatively with GCS score (i.e., their levels decreased gradually with increasing GCS score) and with sampling time (i.e., characterized by a gradual decrease in abundance over time). The metabolites that correlated negatively with GCS score significantly overlapped with those enriched in blood (top 1,000 blood-enriched metabolites; **Fig. 4a, b**) and clear cisternal CSF (**Fig. 4c, d**). This analysis revealed that, among the 598 substances that correlated negatively with sampling time, 278 overlapped with blood-enriched metabolites (46.5%, **Fig. 4a**), whereas only 47 overlapped with CSF-enriched substances (4.5%, **Fig. 4c**). Similarly, among the 279 metabolites that correlated negatively with GCS score, 67 (46.7%) overlapped with blood-enriched substances (11.2%, **Fig. 4b**), whereas a larger number overlapped with CSF-enriched substances (**Fig. 4d**). These findings indicated that some of these negatively correlated metabolites may have originated from blood. These metabolites were gradually excreted via CSF drainage during cisternostomy, leading to a decrease in blood-enriched metabolites in the CSF and contributing to recovery from TBI.

**Fig. 4.**
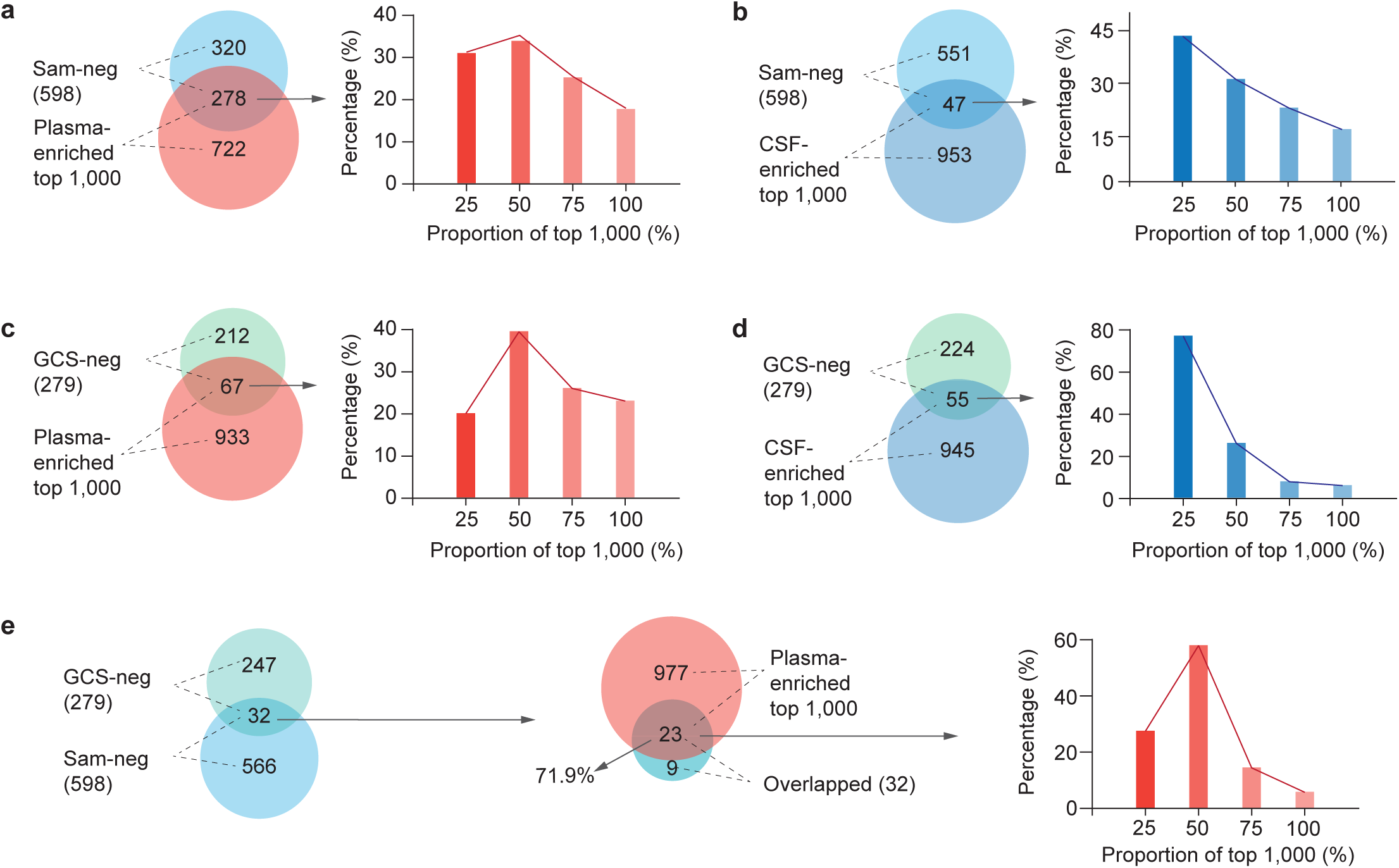
Changes in metabolites in cisternal CSF assessed with analysis after cisternostomy. **a)**, Overlap of 278 metabolites between the *Sam-neg* cluster (metabolites that correlated negatively with sampling day) and the top 1,000 plasma-enriched metabolites. The distribution of these shared features across the 25%, 50%, 75%, and 100% quartiles of the top 1,000 plasma-enriched metabolites was 28.8%, 32.7%, 23%, and 15.5%, respectively. **b),** Overlap of 67 metabolites between the *GCS-neg* cluster (metabolites that correlated negatively with GCS score) and the top 1,000 plasma-enriched metabolites. The distribution of these shared features across the 25%, 50%, 75%, and 100% quartiles of the top 1,000 plasma-enriched metabolites was 17.9%, 37.3%, 23.9%, and 20.9%, respectively. **c),** Overlap of 47 metabolites between the *Sam-neg* cluster and the top 1,000 CSF-enriched metabolites. The distribution of these shared features across the 25%, 50%, 75%, and 100% quartiles of the top 1,000 CSF-enriched metabolites was 40.4%, 27.7%, 19.1%, and 12.8%, respectively. **d),** Overlap of 55 metabolites between the *GCS-neg* cluster and the top 1,000 CSF-enriched metabolites. The distribution of these shared features across the 25%, 50%, 75%, and 100% quartiles of the top 1,000 CSF-enriched metabolites was 72.7%, 21.9%, 3.6%, and 1.8%, respectively. **e),** Overlap of 32 metabolites between the *Sam-neg* and *GCS-neg* clusters. Of these, 71.9% (23 metabolites) originated from the top 1,000 plasma-enriched metabolites. The distribution of these shared metabolites across the 25%, 50%, 75%, and 100% quartiles of the top 1,000 plasma-enriched metabolites was 26.1%, 56.5%, 13.1%, and 4.3%, respectively.

Furthermore, we separately compared the number of features within these overlapping datasets against their relative abundance in blood and CSF. The abundance of enriched substances in blood and cisternal CSF decreased with increasing GCS score and sampling time, and the abundance of substances in the overlaps had an overall downward trend. This suggested that the abundance of certain blood-derived metabolites in the CSF diminished progressively and may have been cleared through the drainage process.

Additionally, we compared the 598 metabolites that correlated negatively with CSF sampling time with the 279 metabolites that correlated negatively with GCS score. We found that 32 metabolites were common to both sets, indicating that the abundance of these substances decreased with both sampling time and the improvement in GCS score. Further analysis of these 32 metabolites revealed that 23 (71.9%) were among the top 1,000 highly enriched compounds in blood. Notably, these metabolites predominantly were within the top 50% in terms of abundance (**Fig. 4e**). These findings suggested that certain negatively correlated substances may have originated from blood clots in AS-CSF following TBI. Moreover, these substances appeared to be progressively eliminated through cisternostomy, facilitated by CSF drainage and the patient’s subsequent recovery.

## Discussion

In this study, we collected a total of 313 cisternal CSF and peripheral venous blood samples from 15 patients undergoing cisternostomy-based CSF drainage treatment for severe TBI. This non-targeted metabolomic study represents the first comprehensive comparison and correlation analysis of the metabolic characteristics of blood and clear cisternal CSF. Our results reveal that blood in the samples was predominantly enriched with lipophilic compounds of higher molecular weight, including phosphatidylcholine, phosphatidylethanolamine, and other glycerophospholipids, whereas cisternal CSF contained a higher proportion of hydrophilic metabolites of smaller molecular weight. Additionally, a significant proportion of metabolites present in CSF during the acute injury phase displayed distinct characteristics compared to those observed in clear CSF during the recovery phase. Notably, many metabolites enriched in acute-stage CSF also were enriched in blood. This suggests that acute-stage CSF contains an abundance of blood-derived metabolites. Their elevated concentrations in the bloodstream likely increase the probability of infiltration into the brain parenchyma via blood clots, resulting in the formation of AS-CSF. Furthermore, some blood-derived metabolites may further accumulate upon entering the CSF.

Using GCS score and the duration of cisternal CSF drainage as key indicators of recovery of TBI patients, we systematically analyzed the alterations in metabolite levels in cisternal CSF during cisternostomy. Metabolites that correlated positively with recovery were predominantly derived from CSF, whereas the negatively correlated metabolites, which decreased in abundance during drainage, tended to be blood-derived. Among the negatively correlated metabolites, 23 blood-enriched compounds gradually decreased in abundance in cisternal CSF as GCS score and sampling time increased. These metabolites, including N8-acetylspermidine and N-acetylputrescine, represent blood-derived compounds consistently excreted during recovery of patients with severe TBI. N8-Acetylspermidine overabundance has been implicated in myocardial cell death, ischemic cardiomyocyte injury, and subsequent cardiac dysfunction^46^. Both N8-acetylspermidine and N-acetylputrescine are classified as polyamines^47, 48^ and have been detected in blood, saliva, urine, and CSF. Polyamines are linked to arginine metabolism and the urea cycle. N-Acetylputrescine, which is primarily derived from amino-acid catabolism, is somewhat cytotoxic and is produced during cadaver decomposition^49^. It has also been suggested as a potential biomarker for Bachman-Bupp syndrome (commonly known as BABS), a condition characterized by global developmental delays, hypotonia, nonspecific physical abnormalities, and behavioral issues, including autism spectrum disorder and attention-deficit hyperactivity disorder. Previous studies have shown that polyamines contribute to cardiac health as well as brain development and function^50^ but also promote aging^51, 52^. Therefore, maintaining polyamine levels within a physiological range is critical, as dysregulation can exacerbate cancer, neurodegenerative diseases, and aging. Elevated polyamine levels are particularly problematic in neurological disorders such as Parkinson’s disease^53^ and epilepsy^54^, in which significantly higher concentrations have been detected in CSF.

Our findings suggest that cisternostomy combined with CSF drainage can effectively clear blood-derived metabolites, contributing to the removal of metabolic waste. However, this study has limitations due to the complex nature of the surgical procedure involved and the challenge of collecting uniform CSF samples from patient cisterns. The inclusion of only a small cohort of patients with severe TBI—despite yielding 313 CSF and blood samples— may have introduced potential biases due to the sample size. Furthermore, the cohort included patients with varying primary etiologies, such as hypertensive cerebral hemorrhage, head trauma, subarachnoid hemorrhage due to aneurysm, and arachnoid cysts. This heterogeneity, coupled with the aforementioned challenge associated with CSF collection, necessitates further studies to compare the efficacy of cisternostomy treatment across diverse etiologies.

Despite the small sample size, our findings highlight the efficacy of cisternal CSF drainage for the clearance of metabolic waste—and specifically of blood-derived substances—in patients with severe TBI of varying etiology. Our results highlight the clinical relevance of cisternostomy for the removal of metabolic waste after TBI. However, the narrow range of GCS score made it challenging to establish robust correlations between metabolite levels and recovery metrics, leading to inconsistencies between GCS score and sampling-time analyses. Further studies should also include more comprehensive and continuous metrics to evaluate the progression of recovery for TBI patients. Nonetheless, our data strongly suggest that cisternostomy aids in the clearance of metabolic waste and supports recovery.

This study utilized non-targeted metabolomics and did not investigate the permeability of the blood-CSF barrier during recovery. Specifically, we lacked data on blood-derived metabolites in healthy subjects or during the very early stages of injury, and thus further investigation warranted. Future research should explore the potential deleterious actions of blood-derived metabolites in the central nervous system, and functional studies could utilize animal models to determine the origins, neuroprotective properties, and potential mechanisms of action of metabolites that increase during recovery.

## Author contributions

W.G. and Y.W. conceived and supervised the project. W.G. and Y.W. designed the experiments. D.Y. conducted the non-targeted metabolomic measurements using LC-MS. Y.W. collected CSF and blood samples from patients, and S.G., T.H., H.W. and Q.L. assisted with blood and CSF sample collection, centrifugation, aliquoting, storage, and transport. D.Y. and W.G. extracted the samples for metabolomic analysis. W.H., L.D., and D.Y. analyzed the metabolomic data, with W.H. primarily focusing on the differentiation analysis of plasma and cisternal CSF and the classification of cisternal CSF into AS-CSF and clear CSF, and D.L. and W.H. performing the correlation analysis. H.W. and W.G. drafted the manuscript, and all authors contributed to discussions, reviewed, and edited the manuscript.

## Acknowledgements

We thank members of both the Ge laboratory at the Chinese Institute for Brain Research (CIBR) and Wang laboratory at Shanxi Bethune Hospital for discussion and suggestions and Dr. B. Samuels for feedback and critical reading of the manuscript. This work was supported by grants from CAMS Innovation Fund for Medical Sciences (CIFMS), 2024-I2M-ZD-012, the STI2030-Major Projects (2022ZD0204700), the Feng Foundation of Biomedical Research, and the National Natural Science Foundation of China (No. 32170964) to W-P.G. as well as the Science and Technology Department of Shanxi Province (202203021221241) to W.Y.

## Data availability

The datasets generated and/or analyzed in the current study are available from the corresponding authors upon reasonable request.

## Supplementary figure legends

**Supplementary Fig. 1.**
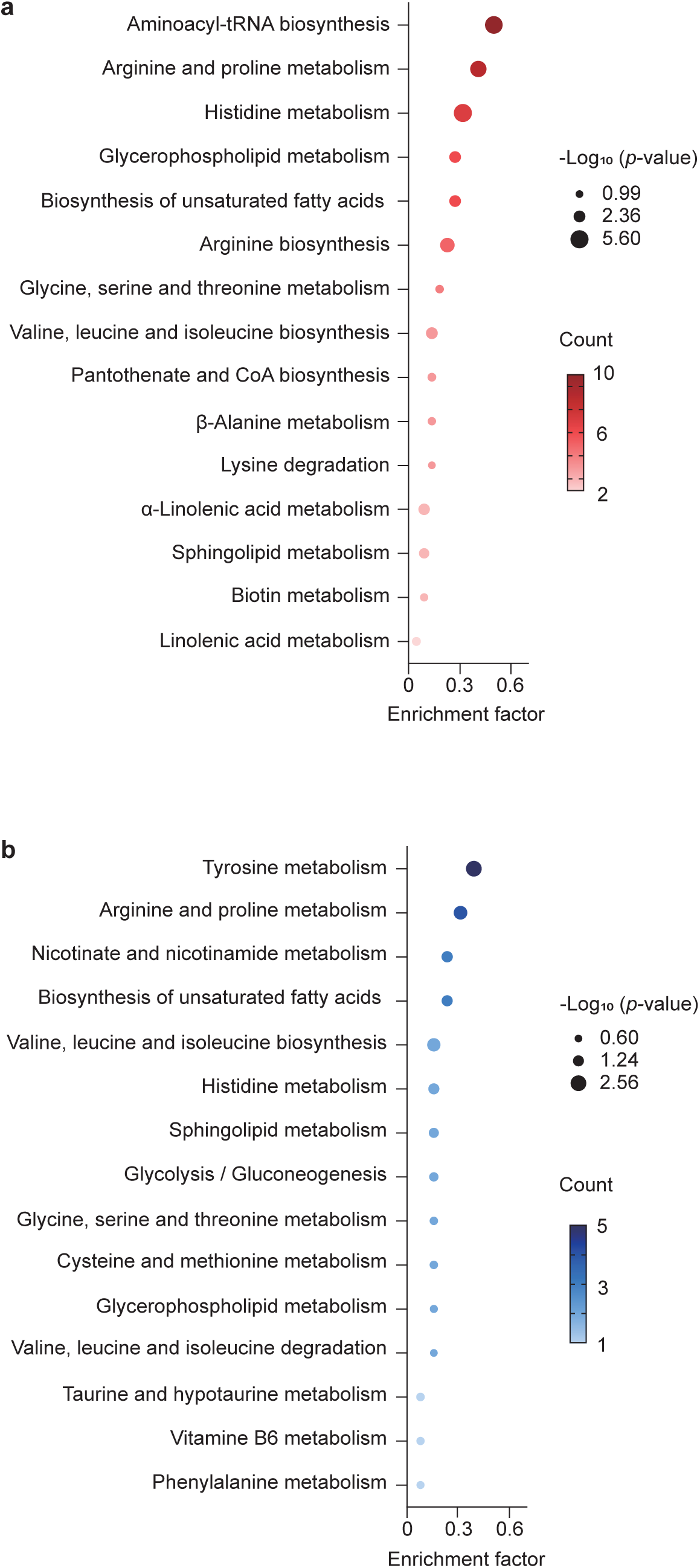
Pathway enrichment analysis of the differential metabolomes between blood and clear cisternal CSF. **a, b)**, KEGG-based qualitative enrichment analysis of metabolites in blood (a) and cisternal CSF (b) to identify the most relevant metabolic pathways. Pathways are ranked by adjusted *p*-values and their pathway impact. Circle size represents pathway impact, whereas color indicates the enrichment significance (*p*-value). KEGG, Kyoto Encyclopedia of Genes and Genomes. LA: linoleic acid. Arg: arginine. Gly: glycine. Ser: serine. Thr: threonine. Ala: alanine. Asp: aspartate. Glu: glutamate. Pro: proline. Lys: lysine. Deg.: degradation. Phe: phenylalanine. Tyr: tyrosine. His: histidine. Cys: cysteine. Tau: taurine. Met: methionine. Vit: vitamin. GPL: glycerophospholipid. Meta: metabolism. Bios: biosynthesis.

**Supplementary Fig. 2.**
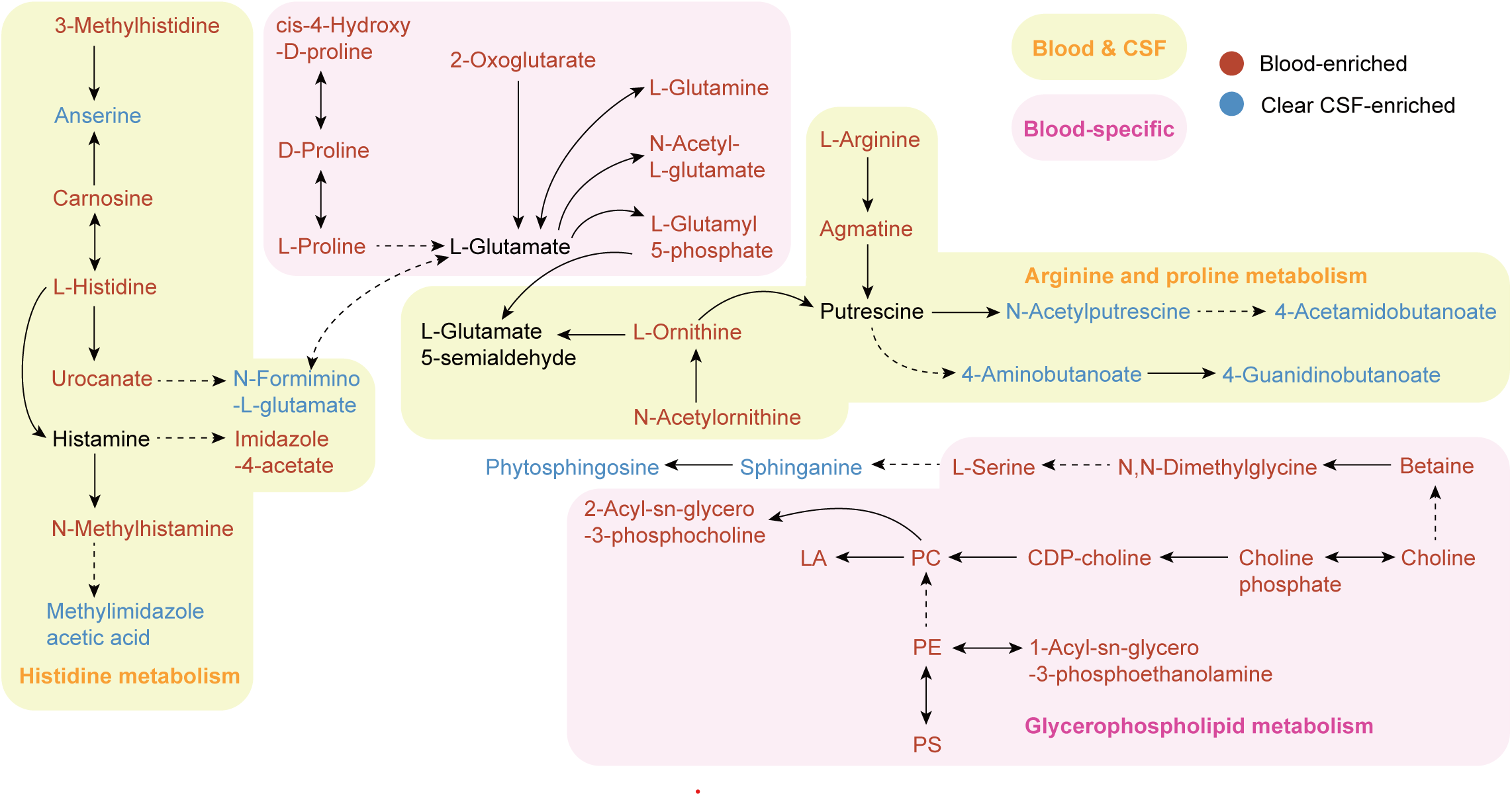
Schematic diagram of pathways enriched in blood and CSF. Schematic diagram illustrating the pathways composed of metabolites enriched exclusively in blood, exclusively in clear cisternal CSF, or shared between both blood and clear cisternal CSF. KEGG: Kyoto Encyclopedia of Genes and Genomes. LA: linoleic acid. Arg: arginine. Gly: glycine. Ser: serine. Thr: threonine. Ala: alanine. Asp: aspartate. Glu: glutamate. Pro: proline. Lys: lysine. Deg: degradation. Phe: phenylalanine. Tyr: tyrosine. His: histidine. Cys: cysteine. Tau: taurine. Met: methionine. Vit: vitamin. GPL: glycerophospholipid. Meta: metabolism. Bios: biosynthesis.

**Supplementary Fig. 3.**
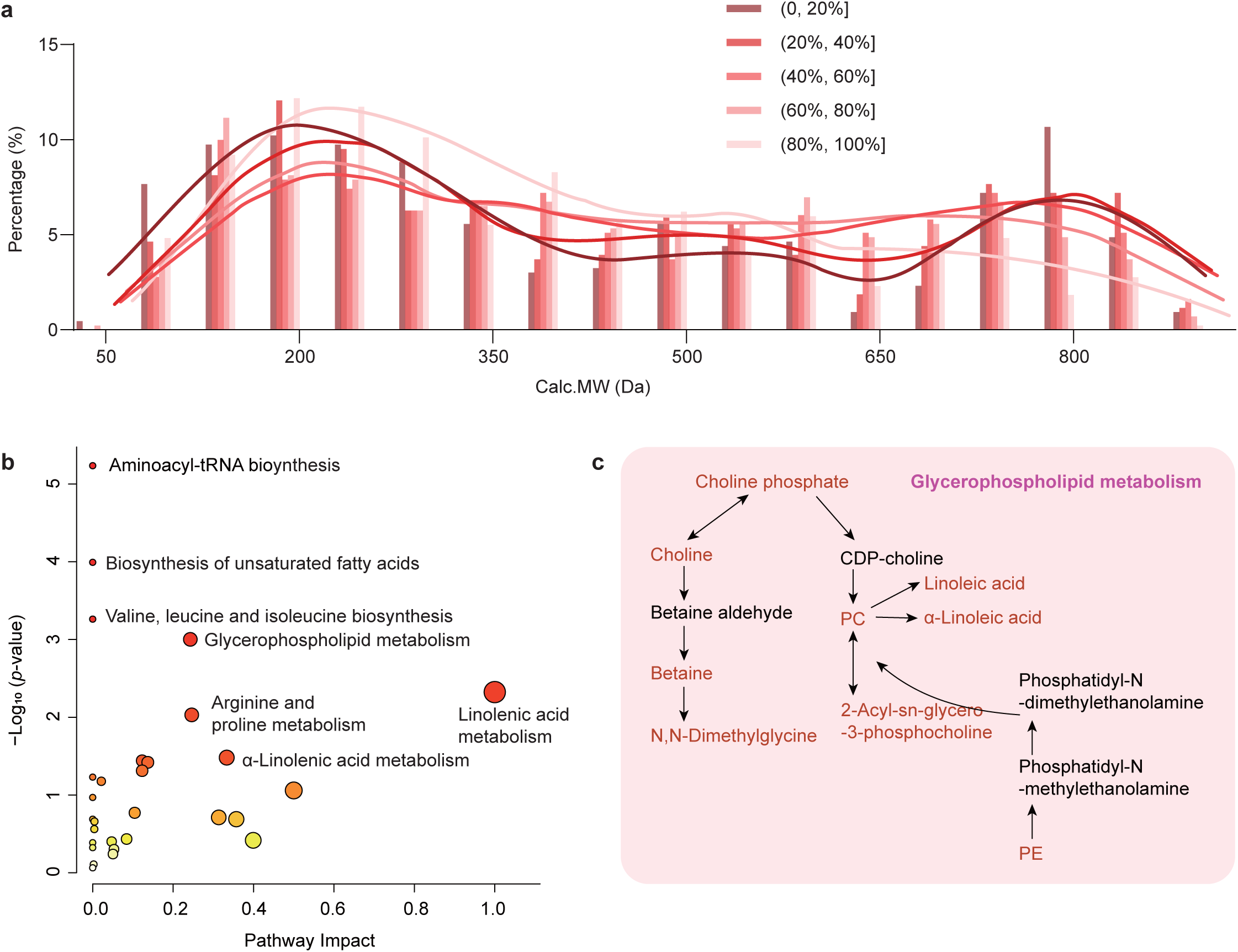
Biochemical pathway–based analysis of metabolites enriched in blood. **a)**, Percentile distribution of blood-enriched metabolites based on molecular mass. The percentiles were calculated by determining the number of metabolites within each 50 m/z segment. **b),** KEGG-based qualitative enrichment analysis of pathways using the top 20% of intensity-based blood-enriched metabolites. Pathways were ranked by adjusted p-values and their impact. Circle size represents pathway impact, whereas color indicates enrichment significance (red indicates higher significance). **c),** Schematic pathway diagrams depicting pathways composed of the top 20% of metabolites enriched in blood. Abbreviations in **(b)** and **(c)**: KEGG, Kyoto Encyclopedia of Genes and Genomes. LA, linoleic acid. Val, valine. Leu, leucine. Ile, isoleucine. Arg, arginine. Pro, proline. Phe, phenylalanine. Tyr, tyrosine. His, histidine. UFA, unsaturated fatty acids. PS, phosphatidylserine. PC, phosphatidylcholine. PE, phosphatidylethanolamine. GPL, glycerophospholipid. meta, metabolism. Bios, biosynthesis.

**Supplementary Fig. 4.**
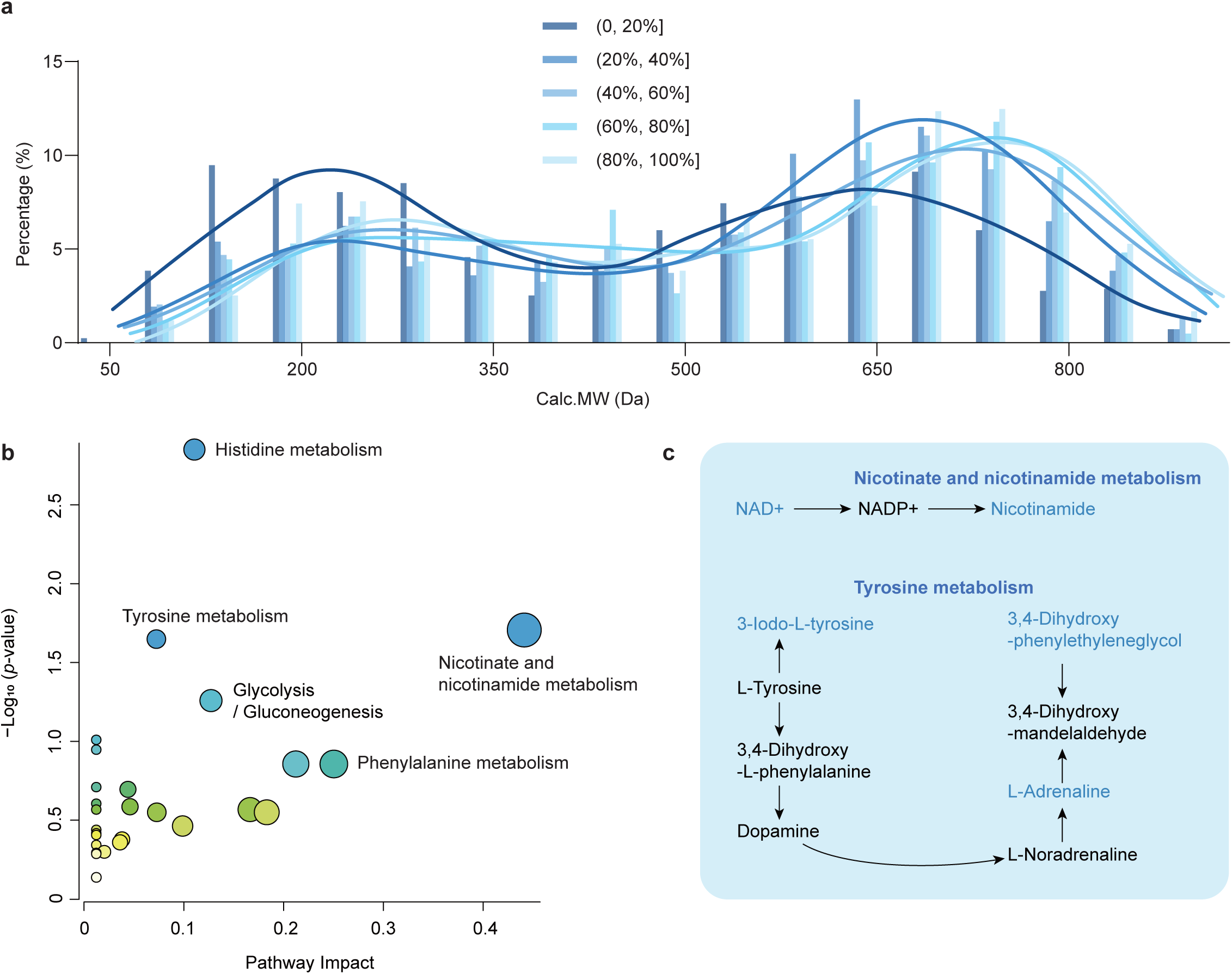
Biochemical pathway–based analysis of metabolites enriched in cisternal CSF. **a)**, Percentile distribution of CSF-enriched metabolites based on molecular mass. The percentiles were calculated by determining the number of metabolites within each 50 m/z segment. **b),** KEGG-based qualitative enrichment analysis of pathways using the top 20% of intensity-based CSF-enriched metabolites. Pathways were ranked by adjusted p-values and their impact. Circle size represents pathway impact, whereas color indicates enrichment significance (red indicates higher significance). **c),** Schematic pathway diagrams depicting pathways composed of the top 20% of metabolites enriched in CSF. Abbreviations in (**a**) and (**b**): KEGG, Kyoto Encyclopedia of Genes and Genomes. LA, linoleic acid. Val, valine. Leu, leucine. Ile, isoleucine. Arg, arginine. Pro, proline. Phe, phenylalanine. Tyr, tyrosine. His, histidine. UFA, unsaturated fatty acids. PS, phosphatidylserine. PC, phosphatidylcholine. PE, phosphatidylethanolamine. GPL, glycerophospholipid. meta, metabolism. Bios, biosynthesis.

**Supplementary Fig. 5.**
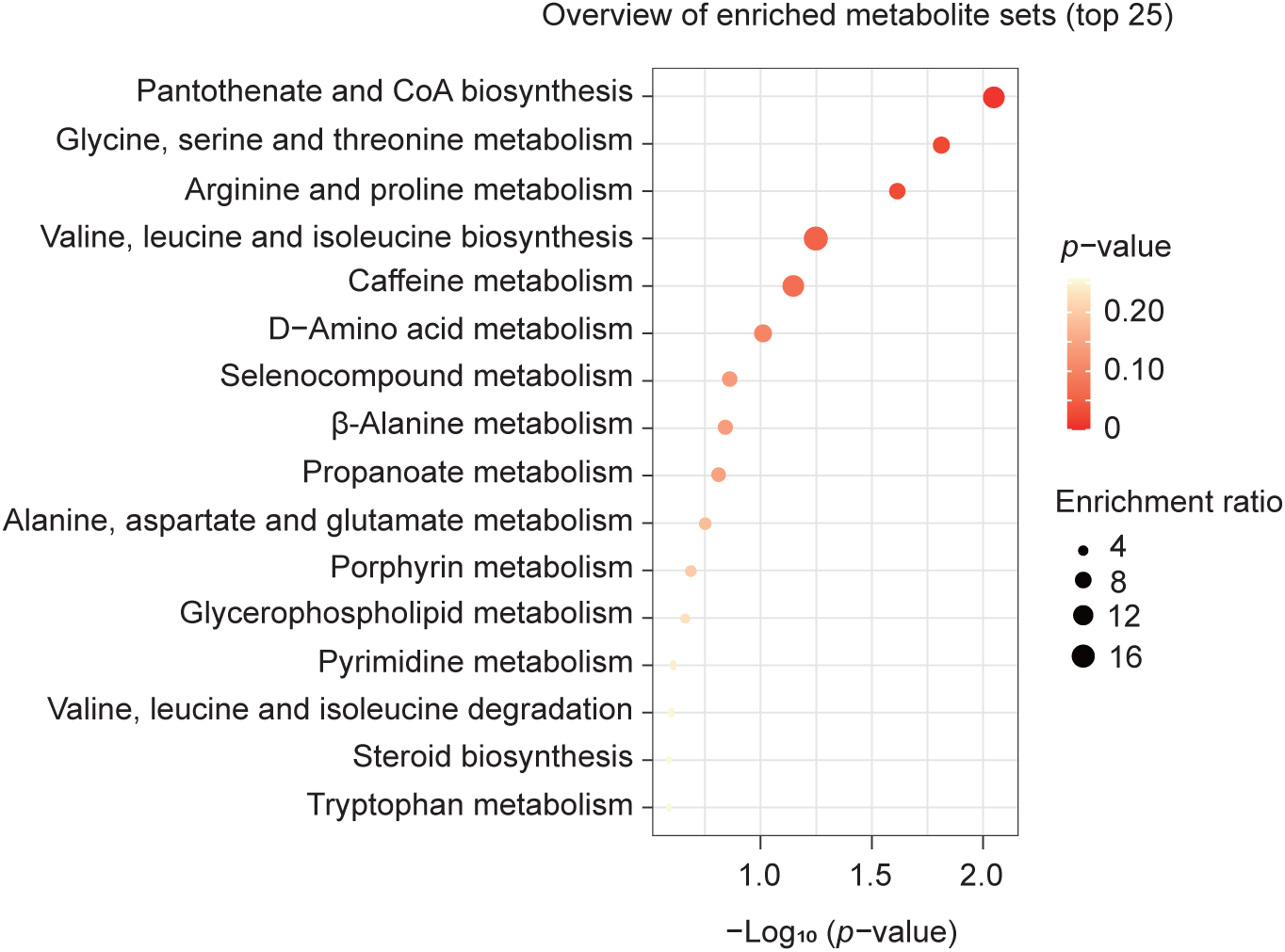
Overlapping metabolites that were enriched in blood but of low abundance in CSF. **a)**, Biochemical pathway–based enrichment analysis of the 918 overlapping features between the AS-CSF and blood samples. The bubble plot shows the top 25 metabolic pathways analyzed from the 918 metabolites. **b),** Circular heatmap displaying the levels of 88 typical metabolic features in blood, AS-CSF, and clear CSF. These metabolites were enriched in blood but of low abundance in clear CSF, and most of them were also enriched in acute-stage CSF. The color bar (in the center) indicates the scale for the levels of standardized metabolites.

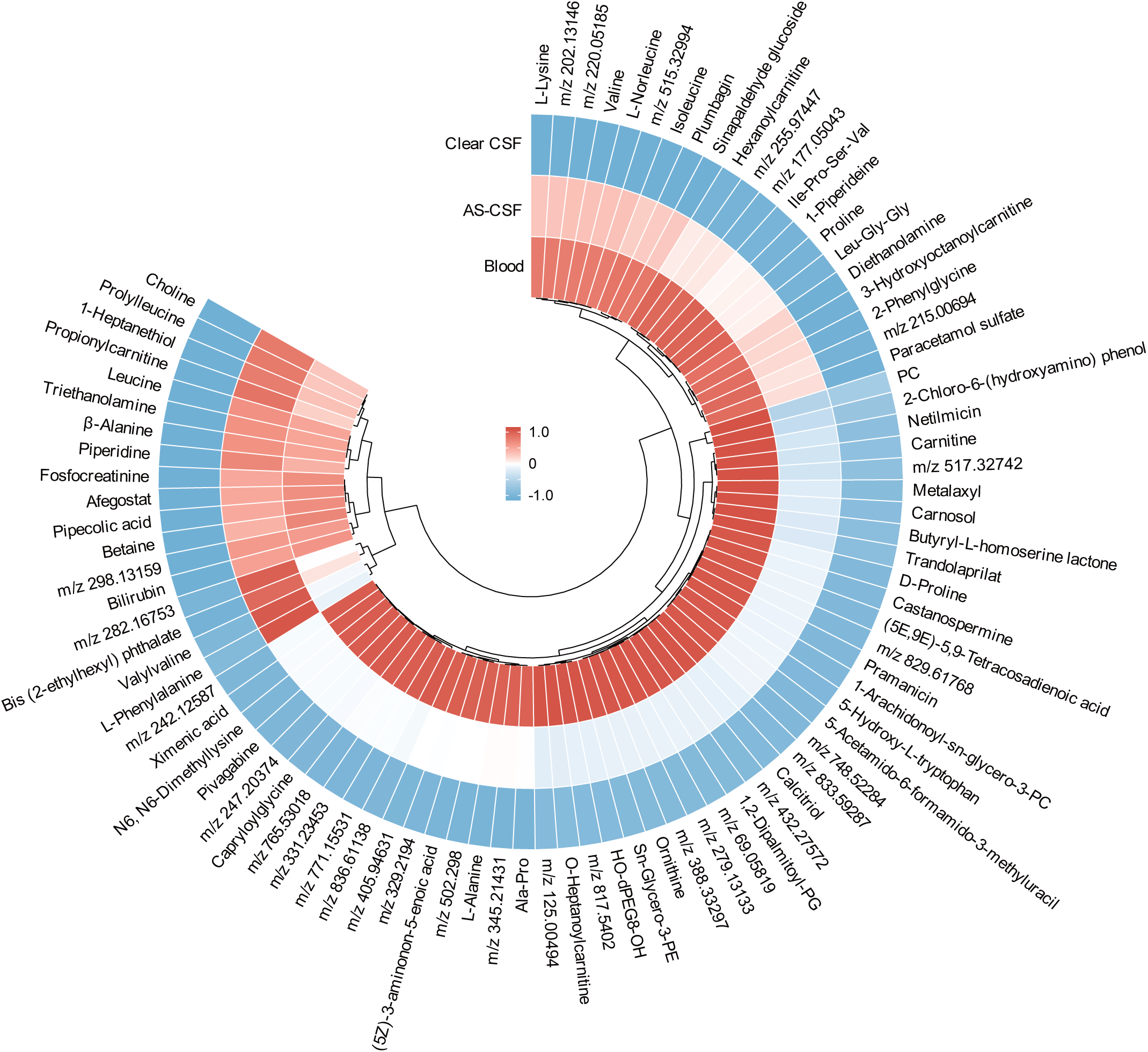

## Methods

### Patient selection

Fifteen TBI patients presenting to the Department of Neurosurgery at Bethune Hospital in Shanxi who required surgical management and met the inclusion criteria were enrolled in the study (Protocol, YXLL-2018-05). All participants signed an informed consent form. The inclusion criteria were as follows: (1) age > 18 years and < 60 years; (2) acute TBI requiring surgery (except for simple epidural hematoma); (3) intracranial pressure > 20 mmHg before craniotomy; (4) no significant changes in vital signs and no surgical contraindications. Exclusion criteria were as follows: (1) concomitant severe underlying systemic illnesses such as abnormal coagulation functions, liver failure, or renal dysfunction; (2) any incidence of severe trauma, penetrating head injury, or preexisting disease; (3) any injury to the thoracic/abdominal cavities. All patients were admitted to the neurosurgical intensive care unit for continuous monitoring of intracranial pressure for one week after surgery, assisted breathing with endotracheal intubation and a ventilator, sedation and analgesia, and symptomatic supportive treatment.

### Classification of cisternal CSF and sample collection information

To evaluate alterations in CSF composition at different stages of the acute phase in TBI patients who underwent cisternostomy, our study classified cisternal CSF samples based on GCS score, timing of sample collection, and CSF clarity. Cisternal CSF from patients with mild to moderate brain injury (GCS score 9–15) was collected at a later stage of recovery and had optimal clarity. In contrast, AS-CSF was associated with a GCS score of 3–8, indicating severe brain injury. These samples were collected earlier in the acute phase, typically clearing by days 1–2 post-initial collection, and had visible bloody turbidity between days 3–6.

Samples that met the predetermined criteria were segregated into distinct groups, with each patient contributing nearly equivalent quantities of samples. The final selection comprised the following: 20 samples of clear cisternal CSF from 5 patients, 16 samples of AS-CSF from 6 patients, and 20 peripheral venous blood plasma samples from 10 patients.

### Collection of peripheral plasma and cisternal CSF

CSF samples were collected from TBI patients via basal cistern drainage every 12 h for a total of 14 times following the completion of surgery. Clinical data, including intracranial pressure and GCS score, were recorded, and peripheral blood samples were drawn from arm veins. Both CSF and blood were stored in sterile, heparin-coated anti-coagulation tubes. The CSF and plasma supernatants were separated by centrifugation at 1,200 x *g* for 15 min at 4°C. All aliquots were stored at –80°C until analysis.

### Metabolite extraction for metabolomic analysis

CSF and plasma samples were thawed at 4°C. Subsequently, 30 μL CSF and 15 μL plasma were transferred into Eppendorf tubes containing 240 μL of ice-cold 90% acetonitrile (ACN)/water (v/v) and 185 μL of ice-cold 86.5% ACN/water (v/v), respectively. Metabolites were extracted, resulting in a final ACN concentration of 80%. After centrifugation at 15,000 x *g* for 15 min at 4°C, 200 μL CSF supernatant and 160 μL plasma supernatant (corresponding to metabolites extracted from 30 μL CSF and 15 μL plasma, respectively) were dried using a centrifugal concentrator (SpeedVac) at room temperature to obtain a pellet. The pellet was subsequently stored at –80°C for metabolomic profiling analysis. Additionally, two types of quality-control pools were prepared by combining equal volumes of CSF and plasma samples, which were then evenly spaced throughout the injection sequence. The samples were re-dissolved in 160 μL of 80% ACN and transferred to a Q Exactive (Thermo Fisher Scientific, San Jose, CA) Orbitrap mass spectrometer for untargeted analysis.

### Non-targeted metabolomic measurement with LC-MS

The metabolites were separated chromatographically using an Agilent Infinity Lab Poroshell 120 HILIC-Z column (2.1 × 100 mm, 2.7 μm) equipped with a PEEK liner and detected using a Q Exactive mass spectrometer. The mobile phases for positive ionization mode were as follows: phase A, water containing 10 mM ammonium formate, and phase B, ACN containing 10 mM ammonium formate; for negative ionization mode, mobile phases were: phase A, water containing 10 mM ammonium acetate, and phase B, ACN containing 10 mM ammonium acetate. The solvent gradient was as follows: 0–4 min, 100% B to 84% B; 4–11 min, 84% B to 40% B; 11–12 min, 40% B; 12–13 min, 40% B to 100% B; and 13–17 min, 100% B. The injection volume was 3 μL, with a flow rate of 0.4 mL/min, and the column temperature was 40°C. All samples were stored at 4°C throughout the analysis period.

Both positive and negative ion scanning modes were employed with the ultraperformance liquid chromatography (UPLC) system to acquire high-quality spectra for the samples, which were interfaced with a Q Exactive Orbitrap mass spectrometer equipped with an electrospray ionization source. For MS1 detection, the automatic gain control target was set to 3,000,000 with a maximum injection time of 100 ms. The resolution was set to 70,000 in full-scan MS mode, with a scan mass range of 60–900 m/z. MS2 detection was performed at a resolution of 17,500, covering a mass range of 60–400 m/z and 350–900 m/z for the mixed quality-control samples. The automatic gain control target was set to 50,000 with a maximum injection time of 80 ms, and the minimum automatic gain control target was set to 1,000, with a dynamic exclusion duration of 6 s.

### Data preprocessing and statistical analysis

The collected LC-MS/MS raw data files were imported into Compound Discoverer 3.2 software (Thermo Fisher Scientific) to obtain matched peak data. A total of 28,244 and 14,637 features were detected in positive and negative ionization modes, respectively. To ensure data quality, data cleaning was performed as follows: First, an initial precursor mass deviation of up to 5 ppm and a fragment mass deviation of up to 20 ppm were allowed. Next, the signal-to-noise ratio was calculated as the ratio of the average intensity of all samples to that of the blank, and it was maintained above 2. Additionally, the average intensity of all samples was required to be above 3 × 10⁴. After applying these screening criteria, 13,114 features were retained.

The peak intensity data, representing the abundance of metabolites in each sample, were normalized using Z-score transformation. The selected compounds were identified using the Human Metabolome Database (HMDB) database and ion fragments obtained from the mass spectrometry analysis. Differential metabolite data were analyzed using one-way analysis of variance with GraphPad Prism 9.0 software. Pathway enrichment analysis was performed through the MetaboAnalyst platform, and related pathways for the differential metabolites were analyzed using the KEGG online database.

